# Understanding Lesion Progression in a Chronic Model of Cerebral Cavernous Malformations through Combined MRI and Histology

**DOI:** 10.1101/2022.11.15.516639

**Authors:** Delaney G. Fisher, Khadijeh A. Sharifi, E. Zeynep Ulutas, Jeyan S. Kumar, M. Yashar S. Kalani, G. Wilson Miller, Richard J. Price, Petr Tvrdik

## Abstract

Cerebral cavernous malformations (CCM), also known as cavernous angiomas, are blood vessel abnormalities comprised of clusters of grossly enlarged and hemorrhage-prone capillaries. The prevalence in the general population, including asymptomatic cases, is estimated to be 0.5%. Some patients develop severe symptoms, including seizures and focal neurologic deficits, while others have no symptoms. The causes of this remarkable presentation heterogeneity within a primarily monogenic disease remain poorly understood. To address this problem, we have established a chronic mouse model of CCM, induced by postnatal ablation of *Krit1* with *Pdgfb*-CreERT. These mice develop CCM lesions gradually over 4-6 months of age throughout of the brain. We examined lesion progression in these mice with T2-weighted 7T MRI protocols. Precise volumetric analysis of individual lesions revealed non-monotonous behavior, with some lesions temporarily growing smaller. However, the cumulative lesional volume invariably increased over time and accelerated after about 3 months. Next, we established a modified protocol for dynamic contrast enhanced (DCE) MR imaging and produced quantitative maps of gadolinium tracer MultiHance in the lesions, indicating a high degree of heterogeneity in lesional permeability. Multivariate comparisons of MRI properties of the lesions with cellular markers for endothelial cells, astrocytes, and microglia revealed that increased cell density surrounding lesions correlates with stability, while increased vasculature within and surrounding lesions may correlate with instability. Our results lay a foundation for better understanding individual lesion properties and provide a comprehensive pre-clinical platform for testing new drug and gene therapies for controlling CCM.

## Introduction

Cerebral cavernous malformations (CCM), also referred to as cavernous angiomas, are hemorrhage-prone, slow flow venous lesions that arise in the central nervous system and affect approximately 0.5% of the general population (Al-Holou et al. 2012; Ene, Kaul, and Kim 2017; Vernooij et al. 2007). CCM form as a result of bi-allelic mutations in one of the three main causative genes—*KRIT1, CCM2*, or *PDCD10* (Snellings et al. 2021). Homozygous germline mutations are embryonic lethal in mice and, presumably, humans (Chan et al. 2011; Kleaveland et al. 2009; Whitehead et al. 2004). Heterozygous germline loss-of-function mutations show variable frequency in different human populations (Akers et al. 2017; Gunel et al. 1996). While usually asymptomatic outwardly, CCM have been associated with increased vascular permeability (Mikati et al. 2015; Yadla et al. 2010). Further, the increased incidence of a second somatic mutation can drive the familial form of the disease, which presents with earlier disease onset and a higher lesion burden, compared to sporadic cases caused solely by biallelic somatic mutations (McDonald et al. 2014; Ren et al. 2021).

One outstanding mystery regarding CCM is the diversity of patient symptom presentation and the degree of symptom severity (Denier et al. 2004; Gault et al. 2006; Gianfrancesco et al. 2007). Some patients of CCM can experience disabling symptoms that commonly include weakness, numbness, severe headaches, vision changes, and difficulty speaking and understanding. More severe symptoms can include stroke, seizures, and even paralysis. Meanwhile, other patients remain asymptomatic. Symptoms typically arise due to cavernomas that have hemorrhaged. However, the trajectory of clinical outcomes of lesions remains unpredictable. Many investigations have been conducted to better understand the heterogeneity of disease severity at the genetic level. For instance, it has recently been discovered that *PIK3CA* gain of function mutations can co-exist with CCM mutations within lesions and may be associated with more severe disease presentation (Hong et al. 2021; Ren et al. 2021; Weng et al. 2021).

Studies have been conducted to utilize MR scans of patients to correlate lesion stability (i.e. growth and hemorrhage risk) with MR-assessed features of permeability and susceptibility (iron deposition) (Girard et al. 2017; Mikati et al. 2014; Tan et al. 2016; Zeineddine et al. 2018). While these studies show correlations of instability with lesion permeability and susceptibility, they fail to elucidate the molecular or cellular basis within and surrounding lesions that correspond with these MR features. Despite the extensive effort to develop predictive models (Girard et al. 2021; Sone et al. 2022), the variability in patient presentation remains to be wholly explained. Recommendation of clinical treatment cannot be fully informed without understanding of lesion trajectory.

Animal models of CCM that reflect the human pathology and patient heterogeneity are necessary for uncovering the heterogeneity seen in patients. Traditionally, acute animal models have been used for rapid screenings of pharmaceutical agents, but they lack many key features of the human pathology. Recently, strides have been made in developing chronic models of the disease that better encompass the human pathology. Cre-inducible models have proven useful for emulating loss of CCM gene heterozygosity in humans. Indeed, chronic models of *Pdcd10* and *Ccm2* mutations have recently been generated and characterized (Cardoso et al. 2020; Detter et al. 2020). *Krit1* mutations are the most prevalent causation of the disease. However, chronic models of *Krit1* have had limited characterization and have been largely confined to postnatal tamoxifen induction at P1, restricting lesions to the cerebellum and retina (DiStefano and Glading 2020; Mleynek et al. 2014).

Despite progress with these new models, most have yet to be robustly characterized with clinically relevant procedures. As MRI is a staple of CCM diagnosis and monitoring in patients, it is equally needed in pre-clinical studies for interpreting mechanistic investigations of CCMs in the context of human disease and for designing improved therapeutic interventions. Prior to therapeutic intervention, it is imperative to understand the baseline progression of the disease without therapeutic intervention and to determine optimal timing of therapeutic intervention.

Furthermore, MR sequences that can evaluate lesion permeability and hemosiderin deposition have been developed in patients with the goal of predicting stability of cavernomas in terms of their growth and hemorrhage (Girard et al. 2017; Hart et al. 2013; Mikati et al. 2015; Mikati et al. 2014). Developing these MR sequences for mouse models of CCM would enable improved assessment of therapeutic intervention risk (i.e., acute hemorrhage due to therapy) and allow for longitudinal monitoring of therapeutic efficacy towards stabilizing the volume and bleeding of treated cavernomas. To this end, we have generated a chronic, tamoxifen inducible *Krit1* model with reduced tamoxifen dose and delayed induction that recapitulates the human CCM pathology. We then developed MRI protocols to enable longitudinal characterization of individual lesion progression throughout the whole brain in terms of lesion volume and permeability. We ensured that our MRI protocols align with immunohistochemical staining of lesions. Further, we identified relationships between our MRI-assessed features of lesions and cellular responses within and surrounding individual lesions. Together, this study lays the foundation for better prognostication of CCM lesion behavior.

## Results

### Delayed postnatal deletion of *Krit1* in the *Pdgfb* domain generates a chronic CCM model with multiple lesions distributed throughout the brain

To study CCM lesion properties in the adult mouse brain, we developed a genetic strategy that gradually generates cavernomas throughout the entire brain over the young adult life span. Traditionally, tamoxifen-induced deletion of conditional CCM alleles was initiated soon after birth (postnatal days P1-P3). However, this early timing severely affects the rapidly developing murine cerebellum, leading to the formation of multiple hemorrhage-prone lesions and a high mortality around one month of age (Detter et al. 2020). We have therefore delayed CCM gene ablation to postnatal day 5-7 (P5-P7). In our approach, we crossed the males of the *Pdgfb* ^*iCreERT2-IRES-EGFP*^ strain (Claxton et al. 2008) (hereafter *Pdgfb-CreERT*) with females harboring the floxed *Krit1* allele (Mleynek et al. 2014). The *Pdgfb-CreERT* studs also carried the null (germline-excised) *Krit1* allele (*Krit1* ^*fl/null*^), emulating familial inheritance pattern of the disease. Only heterozygous progeny of the *Pdgfb-CreERT*; *Krit1* ^*fl/null*^ genotype were used. A single injection of dilute tamoxifen in the dorsal subcutaneous region reliably induced lesion formation beginning at 1 month of age and extending through young adult life (4-6 months; **Figure 1A**). Lesions form throughout the whole brain; frequently in the periventricular striatum and along the hippocampal folds, but are also regularly found in the cerebellum, olfactory bulb, thalamus, cerebral cortex and brainstem (**Figure 1B, Supplemental Figure 1)**. Most mice of this model appear grossly similar to their littermate controls; however, some mutants developed anal prolapse or hydrocephalus. We used a T2-weighted MRI sequence (T2-SPACE described in detail below) to detect lesions in live animals. We confirmed that hypo-intensities detected with T2-SPACE *in vivo* can be unequivocally matched with vascular lesions identified in brain cryosections from these animals with isolectin IB4 (**Figure 1C**), establishing a robust correlation between MRI and histology. Thus, delayed ablation of *Krit1* generates CCM lesions over time throughout the brain and these lesions can be readily identified with MR imaging, and cross-examined with histology and immunohistochemistry.

**Figure 1.**
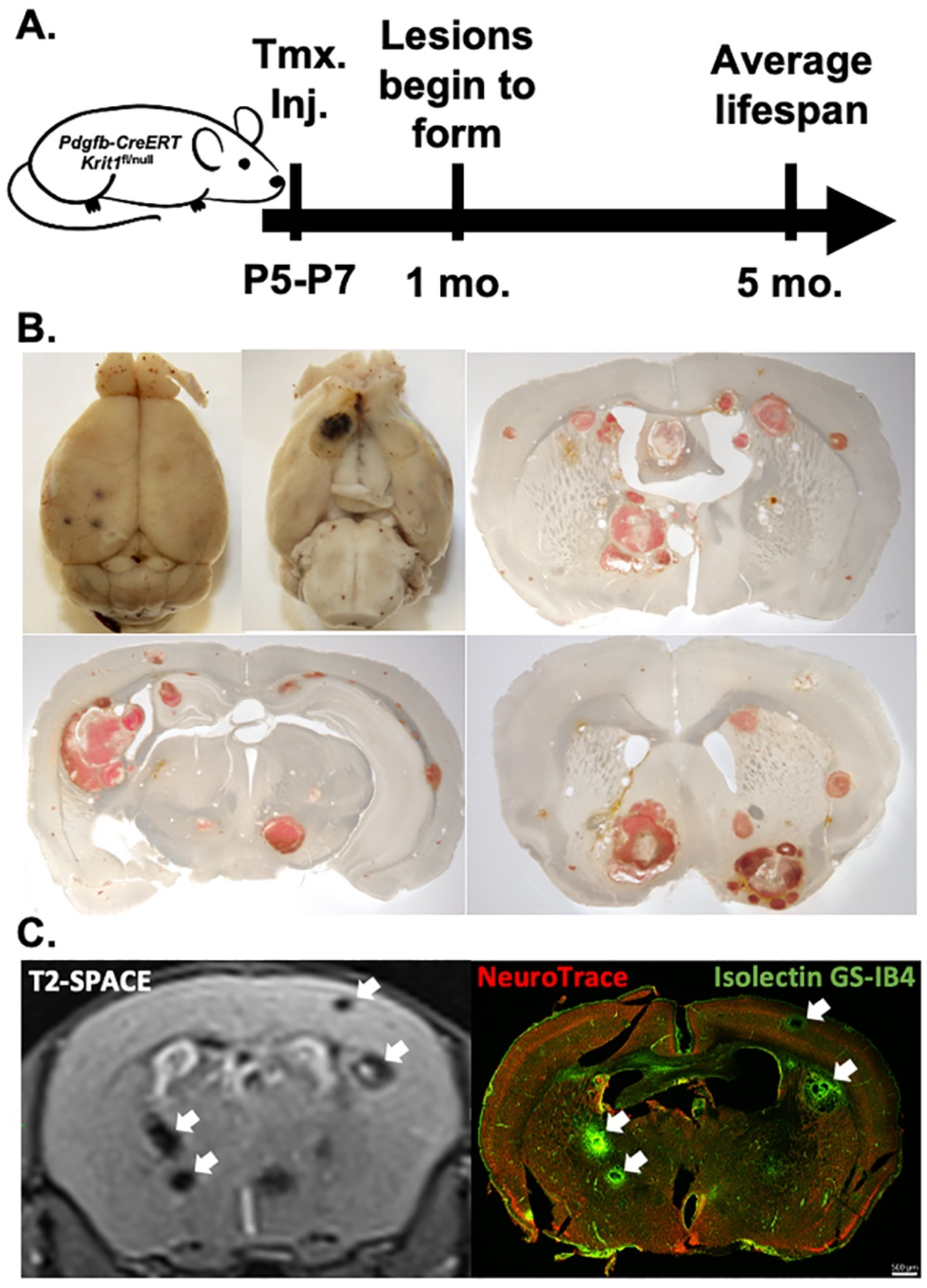
Induction of Krit1 ablation at postnatal day 5 or 7 generates chronic CCM murine model with gradual lesion development brain-wide. (A) Timeline of chronic CCM model generation and disease phenotype. (B) Macroscopic and brightfield images of lesion burden demonstrate lesions form throughout the entire brain. (C) Comparison of MRI (left) and IF image (right) of the same CCM mouse brain section demonstrates alignment of lesions between the two imaging modalities. In IF image, lesions are stained with isolectin GS-IB4 (green) and neurons with NeuroTrace (red). Scale bar=500μm. White arrows denote lesions.

### Longitudinal MRI demonstrates dramatic increases in individual lesion size and total lesion burden with age

We longitudinally characterized lesion burden using T2-weighted SPACE (T2-SPACE) magnetic resonance imaging (MRI). SPACE (‘Sampling Perfection with Application optimized Contrasts using different flip angle Evolution’) is a turbo spin echo sequence which enables 3D image acquisition with a high resolution and relatively short scan times (Mugler III, Kiefer, and Brookeman 2000; Mugler 2014; Mugler et al. 2000). Traditional sequences for CCM detection (i.e. T2*-weighted gradient echo and susceptibility weighted imaging) have an increased sensitivity to lesions but comes at the cost of an enlarged distortion of cavernoma size. To more accurately measure lesion size while still producing high resolution images with clinically acceptable acquisition times, we employed the T2-SPACE sequence at 7T magnetic strength to longitudinally characterize lesion volume in our chronic mouse model. This sequence was previously optimized for the 7T ClinScan MRI Animal Scanner (Bruker Corporation) used in this study. We also confirmed that T2-SPACE closely depicted the lesion dimensions and internal architecture (Liang et al. 1999; Wang, Idowu, and Lin 2017).

Next, we used longitudinal T2-SPACE on 9 mice from 5 distinct litters at 1 month of age (n=3 from 2 litters), 2 months (n=7 from 4 litters), 3 months (n=3 from 2 litters), 4 months (n=5 from 4 litters), and 5 months (n=1; **Figure 2**). This imaging revealed that CCM formation began in low numbers at 1 month of age, and CCM formation progressed with age throughout the entire brain. By tracking the total lesion number and combined lesion volume for a given mouse at each imaging timepoint (**Supplemental Movie 1**, [link](https://drive.google.com/file/d/1ckbTO14iKgTaz1mB-RVaW6molOUVGhMS/view?usp=share_link)), we found that cumulative lesion burden increases with age and accelerates after 3 months of age (**Figure 2A-B**). The median total lesion volume in the brain for a mouse was 0.041mm^3^ at 1 month of age, 0.044mm^3^ at 2 months, 0.714mm^3^ at 3 months, 18.486 mm^3^ at 4 months, and 46.721 mm^3^ at 5 months. T2-SPACE images of each mouse were then analyzed to track individual lesions across imaging timepoints. This individual lesion analysis revealed that lesions in every mouse displayed variable growth rates, including negative rates (i.e. shrinkage of lesion size; **Figure 2C**). However, the majority of individual lesions increased in size with age. Altogether, our chronic *Krit1* CCM mouse model displays progressive lesion formation and increasing cumulative lesion burden with age, which is seen in patients with familial CCM (Denier et al. 2004; Labauge et al. 1998; Rigamonti et al. 1988). These results indicate that our model is well-suited for studying individual and variable lesion dynamics with context relevant to the human pathology.

**Figure 2.**
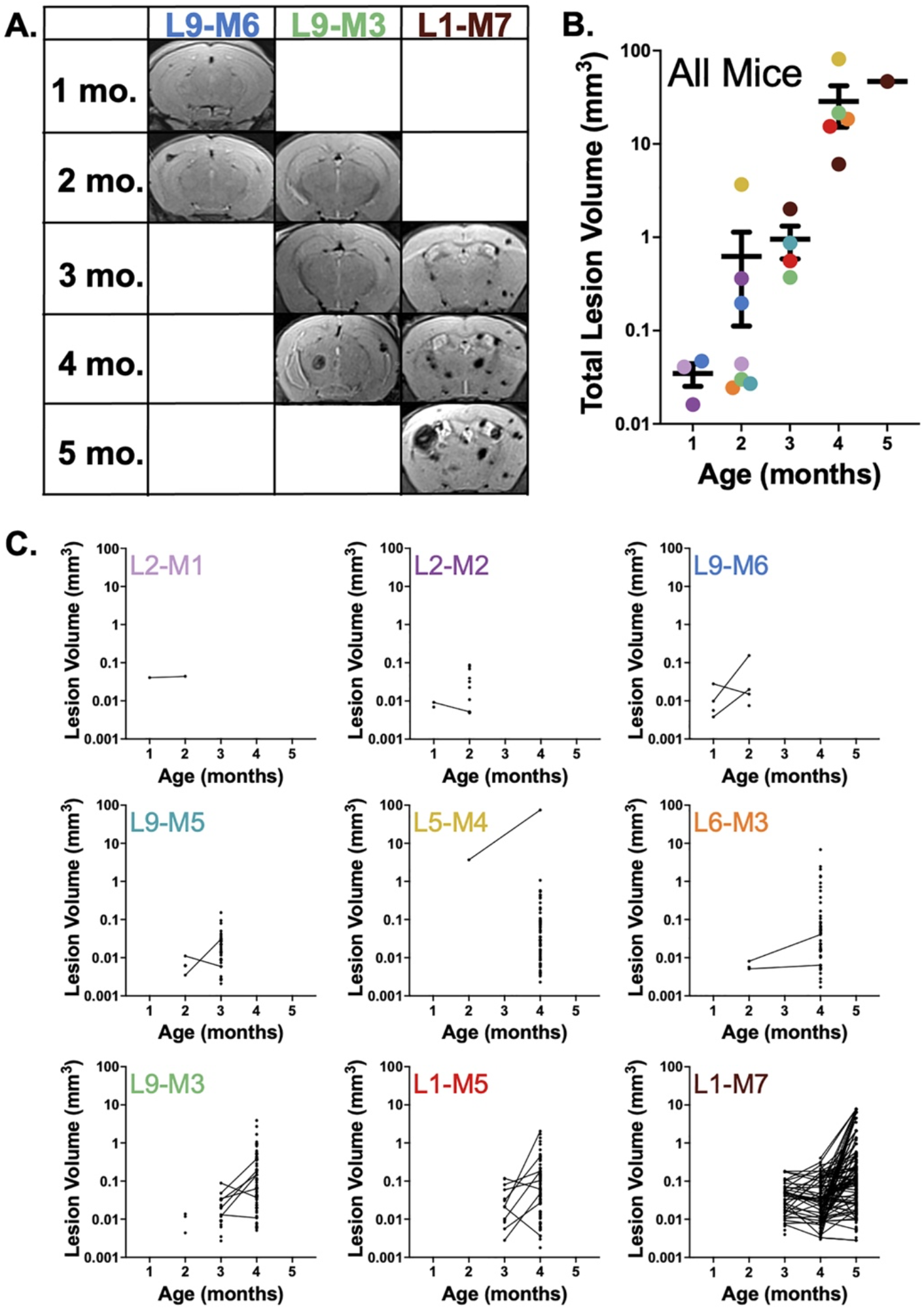
Volumetric analysis of lesions from longitudinal T2-SPACE MR images reveals dramatic increases in lesion burden with age and dynamic changes in size of individual lesions. (A) Representative T2-SPACE MR images from 3 mice in our cohort, illustrating formation of new lesions and dynamic changes in lesion size across time points. (B) Graph of combined lesion volume within individual mouse brains as a function of age for all mice in our cohort. Individual mice represented as a single colored dot, which corresponds with graph title color in panel C. (C) Graphs of lesion volume as a function of age for each mouse in our cohort. Lines indicate lesions identified as the same lesion across imaging time points. Dots without lines indicate de novo lesion formation or lesions that were only identified at a single time point. Graph titles indicate unique mouse ID where L# denotes litter number and M# denotes arbitrary mouse number within litter.

Of note, susceptibility-weighted imaging (SWI) MRI sequences are also used due to their increased sensitivity to inhomogeneities in magnetic field strength surrounding iron-rich lesions. This sequence often produces what is called a “blooming” artifact (Awad and Polster 2019; Flemming and Lanzino 2020). We tested a SWI sequence optimized for the 7T ClinScan MRI scanner and imaged CCM animals immediately prior and after acute LPS-mediated inflammation. This approach demonstrated a greater sensitivity to post-infection lesion increases due to presumed bleeding (**Supplemental Figure 2**). However, this increased sensitivity to susceptibility effects also enlarges lesions in the produced image by distorting their perceived size as previously described (Greenberg et al. 2009; Hudnall et al. 2021), disqualifying this approach from accurate volumetric assessment of CCM lesions.

### T1 Contrast Maps of MultiHance deposition reveal individual lesion permeability

We next sought to characterize the lesion “leakiness”, or internal permeability, with MultiHance, a ∼1 kDa MRI gadolinium-based contrast agent. We used a modified version of dynamic contrast enhanced (DCE) MRI (Tofts and Kermode 1991) that we have termed T1 contrast mapping, as described in the Methods. Briefly, T1 maps were constructed from MR images acquired from CCM mice prior to and following a MultiHance bolus injection. The T1 changes due to contrast accumulation were used to determine local concentrations of the contrast agent. In practice, the total deposition of MultiHance serves as a measurement of lesion leakiness, which lends insight into the potential hemorrhage risk of a lesion and its clinical instability. Compared to traditional DCE methods, which provide temporal information about gadolinium accumulation but are typically confined to a few MR slices of the brain (Hylton 2006), our T1 contrast mapping approach enables whole brain analysis of lesion permeability.

We employed longitudinal T1 contrast mapping on 6 of the 9 mice that received volumetric image analysis. A dataset was acquired for ages of 1 month (n=1), 2 months (n=4 from 3 litters), 3 months (n=2 from 1 litter), 4 months (n=3 from 3 litters), and 5 months (n=1; **Figure 3**). T1 mapping generated gadolinium concentration maps that were aligned with T2-SPACE sequences, enabling us to quantify MultiHance deposition in T2-SPACE-defined lesions (**Figure 3A; Supplemental Movie 2**, [link](https://drive.google.com/file/d/1dRJjiDYenf3XhZzGN5XFEUjM9ucAtYxi/view), **and 3**, [link](https://drive.google.com/file/d/1oPM9OcCzWgQYrLnPchPYg88TlK4QqI_M/view)). By tracking the combined lesion gadolinium deposition for a given mouse at each imaging timepoint, we found that cumulative lesion leakiness increases with age and appears to accelerate after 4 months of age (**Figure 3B**). Median total gadolinium deposition was 3.26 *μ*g at 1 month of age, 9.41 *μ*g at 2 months, 31.7 *μ*g at 3 months, 533 *μ*g at 4 months, and 1,600 *μ*g at 5 months. The gadolinium concentration maps of each mouse were then analyzed to track individual lesions across imaging timepoints. This individual lesion analysis revealed that all T1-mapped lesions displayed variations in gadolinium deposition over time, with some lesions even displaying decreased gadolinium deposition at later time points (**Figure 3C**). However, the majority of individually tracked lesions increased in gadolinium deposition with age. Taken together, our data shows that the chronic *Krit1* CCM mouse model is characterized by progressive lesion burden with cumulative increases in lesion number, volume, and leakiness with age. Individual lesions can demonstrate variable changes in size and leakiness, which has also been in seen in human patients (Hart et al. 2013; Labauge et al. 2007; Mikati et al. 2015; Morrison et al. 2003; Pozzati et al. 1996).

**Figure 3.**
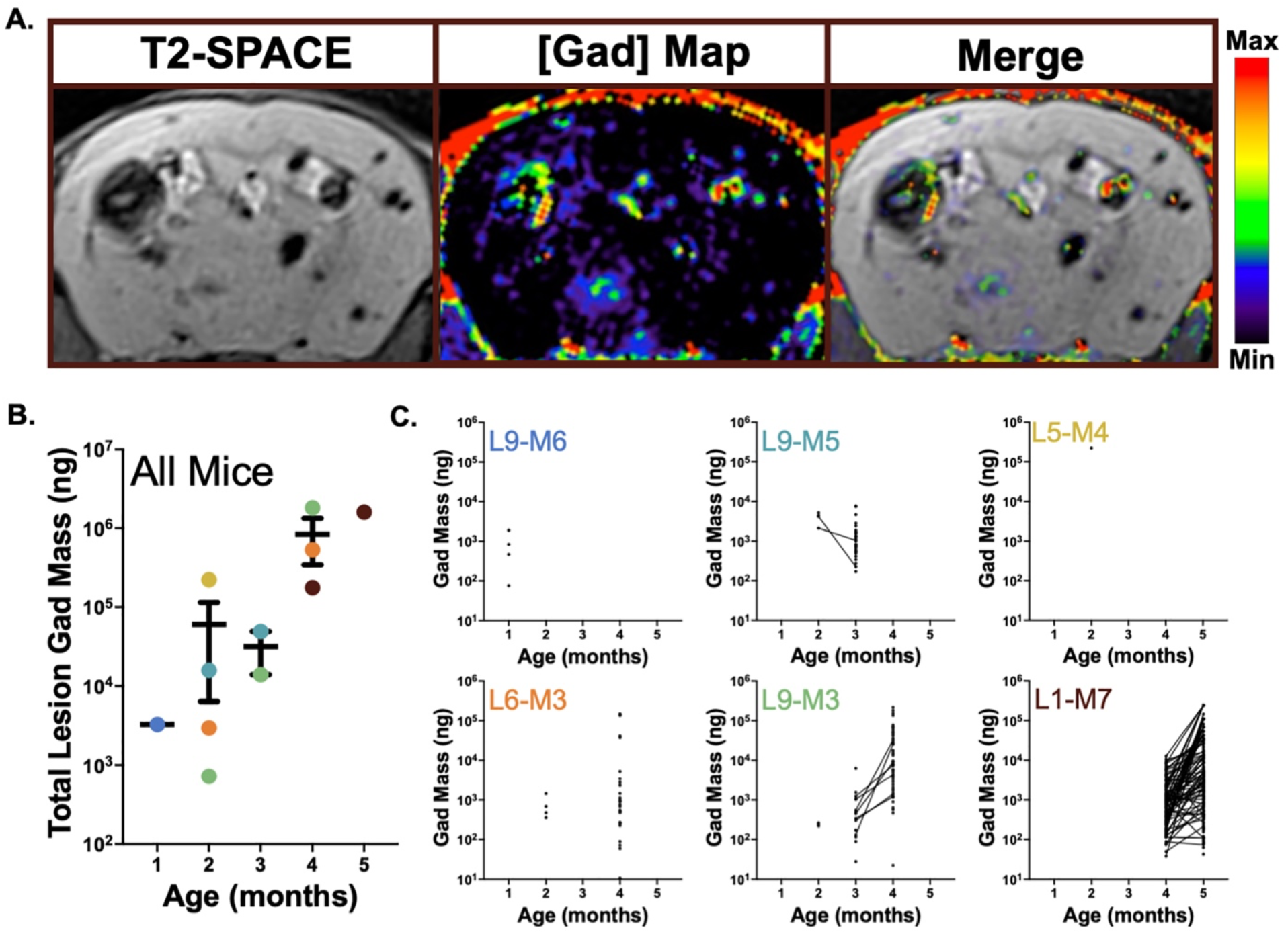
Permeability analysis of lesions from longitudinal T1 contrast mapping reveals cumulative increase in lesion permeability with age while individual lesion permeability varies over time. (A) Representative MR images of T2-SPACE, gadolinium concentration map (generated from T1 contrast mapping), and merged image of the same brain section, illustrating leakiness of individual lesions in terms of gadolinium deposition. In the gadolinium concentration map, hyperintense areas indicate regions with higher gadolinium deposition. (B) Graph of combined lesion gadolinium deposition in mass within individual mouse brains as a function of age for all mice in our cohort. Individual mice represented as a single colored dot, which corresponds with color of graph title in panel C. (C) Graphs of gadolinium deposition in individual lesions as a function of age for individual mice in our cohort. Lines indicate lesions identified as the same lesion across imaging time points. Dots without lines indicate de novo lesion formation or lesions that were only identified at a single time point. Graph titles indicate unique mouse ID where L# denotes litter number and M# denotes arbitrary mouse number within litter.

### CCM lesion permeability is highly variable and correlates poorly with lesion volume

We asked if any correlation existed between lesion volume and lesion leakiness. The total MultiHance deposition in individual lesions was scaled by lesion volume to produce specific gadolinium concentration. Gadolinium concentrations in individual lesions were then plotted against the lesion volume for each imaging point. Concentration versus volume plots were generated for 1 month of age (n=1), 2 months (n=4 from 3 litters), 3 months (n=2 from 1 litter), 4 months (n=3 from 3 litters), and 5 months (n=1; **Figure 4**). A linear regression fitted to these plots revealed no correlation between lesion volume and MultiHance concentration in our measurements. The coefficient of determination (R^2^) values were considerably low (show range of values) for all imaging points, suggesting a poor linear relationship. In other words, our analysis reveals a high degree of heterogeneity of lesion permeability between individual animals, and over their lifespan. This heterogeneity was also observed in lesions within individual mice harboring a sufficient number of datapoints (**Supplemental Figure 3, 4 and 5)**. It is noteworthy that heterogeneity of lesion permeability was also observed in human patients using similar MR techniques (Hart et al. 2013; Mikati et al. 2015). Thus, our mouse model of CCM appears to recapitulate the heterogenous lesion permeability properties seen in human CCM patients.

**Figure 4.**
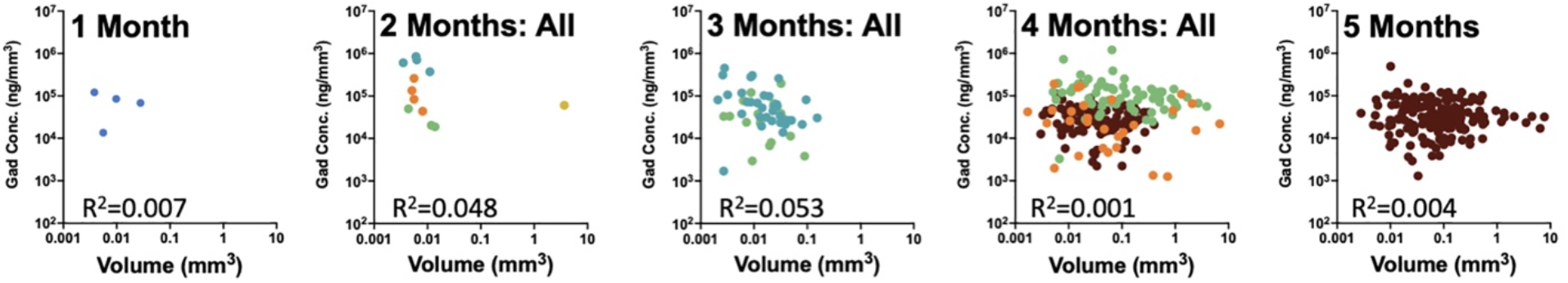
Lesion permeability displays a high degree of heterogeneity. Graphs of gadolinium concentration as a function of lesion volume for individual lesions at each imaging timepoint. Individual mice represented as a single-colored dot, which corresponds with color-coding in Figures 2 and 3. Coefficient of determination values indicates poor correlation between gadolinium concentration and volume for each time point, suggesting highly heterogenous permeability of lesions across age in our chronic CCM model.

### Increased MultiHance deposition in the lesions correlates with decreased cell density in the surrounding parenchyma

Lastly, we asked if the MRI features identified in CCM lesions, namely the lesion volume and lesion permeability, correlated with histological features in close proximity of the lesions. For this analysis, we entirely sectioned the L9-M3 brain and stained with antibodies directed against Iba1 (for microglia), GFAP (for astrocytes) and CD31 (aka PECAM-1, for endothelial cells). The nuclear stain DAPI was included in the mounting media. In total, we identified 17 lesions in confocal montages of the coronal sections that were unequivocally matched with the T2-SPACE MRI sequence recorded immediately prior to brain extraction. High resolution images of the lesions were then used to measure fluorescence signal mean intensity of these three cell populations within the lesion, within the 50-µm perimeter of the lesion, as well as within the 100-µm perimeter of the lesion (**Figure 5, Supplemental Table 1**). The specific cell type intensities were then correlated to lesion volume and MultiHance concentration using the non-parametric Spearman’s rank correlation test. This analysis revealed that cell density outside the lesions, determined as DNA content by DAPI fluorescence within the immediate 50-µm perimeter of the lesion, was inversely correlated with MultiHance concentration inside the lesion (ρ = - 0.61; p < 0.01). The 100-µm ring was also inversely correlated, although at a lower significance level (ρ = - 0.51; p < 0.05) (**Supplemental Figure 6 and 7**). Principal component analysis (PCA) corroborated inverse correlation between MultiHance and DAPI, along with weaker trends towards inverse correlations of MultiHance with microglial and astrocytic densities outside the leaky lesions (**Supplemental Figure 8**). Conversely, the glial density inside the lesion correlated positively, albeit weakly, with MultiHance concentration. The endothelial cell density both inside and outside the lesion showed weak positive correlations with gadolinium concentration. On the other hand, lesion volume did not correlate with MultiHance deposition (ρ = - 0.14, p = 0.58), in line with previous measurements. Of other noticeable trends, microglia and astrocytes both inside and outside the lesions appear to be inversely correlated with lesion volume, whereas endothelial cells both inside and outside of lesions trended positively with lesion volume (**Supplemental Figure 7 and 8**).

**Figure 5.**
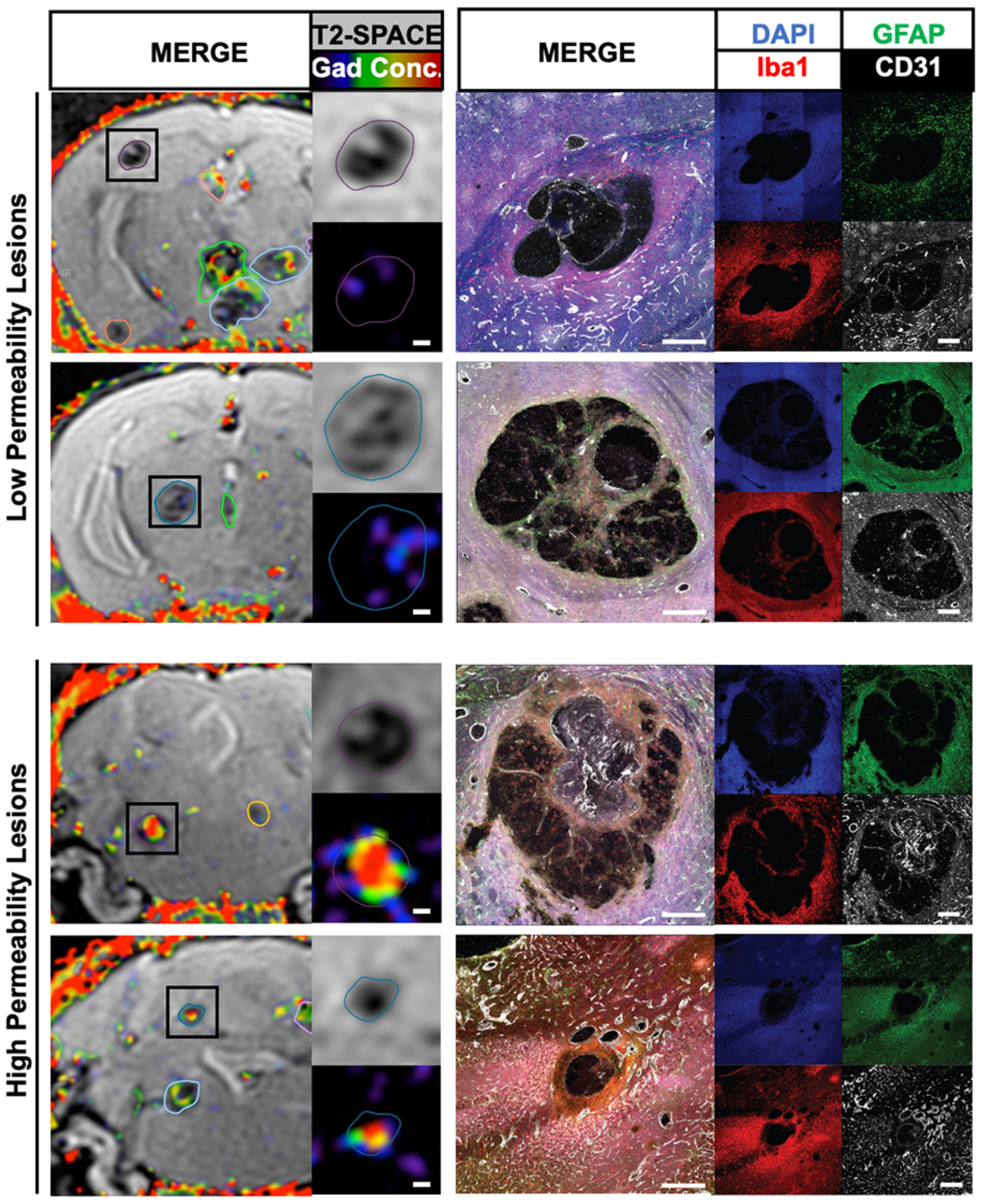
The cellular architecture of CCM lesions with low and high permeability. Two lesions with the lowest permeability (#32 in the subcortical corpus callosum, MultiHance concentration 24.8 μg/mm^3^; and #34 in the thalamus, MultiHance concentration 35.6 μg/mm^3^) are shown on top. Two lesions with the highest permeability (#43 in the brainstem, MultiHance concentration 112.4 μg/mm^3^; and #47 in the cerebellum, MultiHance concentration 151.5 μg/mm^3^) are shown at the bottom. MRI insets show T2-SPACE images and gadolinium concentration maps of each lesion in detail. Matching IHC images of each DAPI, GFAP, Iba1 and CD31 channels are shown on the right, next to the 4-channel overlay. Large, highly permeable lesions typically contain numerous CD31-positive endothelial cells. All scale bars are 200 μm.

Our data points to detrimental effects of high permeability on glial cell density in the parenchyma surrounding the lesions, and suggests that dense vascularization, both inside and outside of the lesions, may have negative effects on lesion stability.

## Discussion

Despite years of investigation, it is still poorly understood why CCM patients have heterogenous presentation in symptoms and lesion phenotypes. Due to the difficulties associated with human studies and limited access to human tissue samples, generation of animal models that reliably capture the human pathology are necessary for studying and furthering treatment of this disease. In this manuscript, we describe a chronic *Krit1* CCM model with delayed tamoxifen induction that replicates human features of lesion development throughout the whole brain and resembles lesion dynamics found in human patients. We also characterize, for the first time, individual CCM lesion progression through the combination of multiple clinical MRI sequences performed longitudinally and immunohistochemical staining of cell populations. This unique merging of a clinically relevant mouse model with advanced MR imaging tools represents a powerful new platform for testing mechanisms of CCM disease, as well as for identifying and pre-clinically testing promising new therapies.

Following generation of our model, we employed clinically relevant MRI sequences longitudinally to characterize lesion progression and dynamics. MRI is commonly used to diagnose and monitor CCM in patients. Despite its ubiquity in the clinic, MR imaging has been largely neglected in many pre-clinical studies, with little data existing on MRI-based characterization of CCM models to robustly assess lesion progression longitudinally. Using T2-SPACE to accurately assess lesion volume and T1 contrast mapping to assess lesion “leakiness,” we find that lesion volume and permeability cumulatively increase with age, while rates of growth and leakiness are variable across individual lesions and time points. Notably, this variability in lesion growth rate and permeability has also been shown in patients (Hart et al. 2013; Labauge et al. 2007; Mikati et al. 2015; Morrison et al. 2003; Pozzati et al. 1996). Our results differ from that in Mikati et al. where patients with familial CCM were shown to not have a correlation between permeability and age (Mikati et al. 2015). However, due to the restriction of their DCEQP protocol (which is confined to 4-6 MRI slices) only a limited number of lesions could be tracked in individuals, whereas our T1 contrast mapping protocol allows for whole brain assessment of lesion leakiness. Thus, we believe that lesion permeability is likely correlated with age for the familial disease if all lesions were examined (Mikati et al. 2015). The development and translation of these clinical MRI protocols for mouse models of CCM has enabled the establishment of the baseline progression of the disease without therapeutic interventions and for assessment of optimal intervention timing in our chronic model. This understanding of how CCMs progress in mice is needed for testing therapeutic approaches. Additionally, progression of CCMs is clinically relevant as unstable lesions (i.e. lesions that increase in size and bleeding) lead to disabling symptoms for patients.

In the clinic, lesion permeability is correlated with increased risk to the patient (Mikati et al. 2015), but relationships that influence lesion permeability remain unknown (Hart et al. 2013). We sought to first identify if lesion volume correlated with gadolinium concentration within lesions in our model. We observed that lesion volume and lesion gadolinium concentration have a poor linear correlation. This finding indicates that there is a high degree of heterogeneity of leakiness in individual lesions throughout individual mice and across mice in our model that is not explained by the lesion’s volume. Similarly, poor correlation between lesion volume and permeability in patients has also been reported (Hart et al. 2013; Mikati et al. 2015).

To further understand lesion heterogeneity in our model, we sought to associate our MR features of lesion volume and lesion leakiness with cellular responses within and around lesions. A limited number of lesions have been correlated from MRI and H&E staining previously (Shenkar et al. 2008), but this is the first association of MRI and specific cell population markers to our knowledge. Spearman non-parametric correlation of our MR features and cell population markers suggested that more permeable lesions contain greater endothelial cell populations within and around cavernomas than less permeable lesions. Meanwhile, less permeable lesions trend toward higher astrocyte and microglial populations surrounding cavernomas than more permeable lesions.

## Conclusion

As chronic models of CCM are designed to test mechanistic and therapeutic avenues that inform clinical practice, it is imperative that these models’ representation of the clinical pathogenesis is validated. To enable clinically analogous validation of our model, we optimized and employed clinical MR protocols. We show that several characteristics of the human pathology is recapitulated in our chronic model of CCM, including: progressive lesion formation throughout the whole brain that increases with age, changes in size and leakiness of individual lesions over time, and heterogenous permeability of individual lesions. We next used the lesion features assessed from MRI (i.e. lesion volume and leakiness) to lend insight to the cellular populations within and surrounding lesions as determined from IHC. We find that increased cell density surrounding the lesion is correlated with lesion stability, while increases in endothelial cells within and surrounding lesions may correlate with lesion instability. This study is the first to establish the baseline conditions of individual lesions in a chronic *Krit1* CCM murine model, providing insight that is essential to advancing treatment strategies for this debilitating disease.

## Materials and Methods

### Animals and Treatments

All animal experiments were approved by the Animal Care and Use Committee at the University of Virginia. The animals were housed under standard laboratory conditions (22°C and 12h/12h light/dark cycle). The *Pdgfb* ^*iCreERT2-IRES-EGFP*^ (*Pdgfb-CreERT*) line was described previously (Claxton et al. 2008). To generate experimental animals, male *Pdgfb-CreERT* mice were crossed to the floxed and null *Krit1* alleles (*Krit1*^*fl*/null^) (Mleynek et al. 2014) to produce the desired genotype *Pdgfb-CreERT*; *Krit1* ^*fl/null*^ males or females. Genotyping was performed by Transnetyx (Cordova, TN) using real-time PCR assays specific for *Krit1* wt, floxed and null alleles, as well as the codon-improved Cre recombinase in *Pdgfb-CreERT. Krit1* gene ablation was induced with a single subcutaneous injection of 50uL of tamoxifen dissolved in corn oil at a concentration of 2mg/mL between postnatal day 5 and 7. To induce inflammatory responses, 0.1 mg/mL solution of lipopolysaccharides in PBS (LPS, Sigma-Aldrich L4391) was injected in 50 µL i.p., corresponding to 0.25 mg/kg dose, 12 h prior to imaging.

### MR Imaging

A 7T small animal MRI scanner (Bruker/Siemens ClinScan) was used to acquire T2-SPACE and T1 contrast images. Mice were imaged monthly, starting as early as one month of age and as late as 5 months of age. To be included in this study, each animal had to have at least two imaging time points conducted. Three-dimensional T2-SPACE MRIs were acquired for all mice in this study using a repetition time of 3000 ms, echo time of 80 ms, pixel size of 125 μm x 125 μm x 100 μm, and 2 averages. Scan time for the T2-SPACE sequence was ∼20 min. T1 contrast mapping was executed by performing 3D spoiled gradient echo sequences at various flip angles before, and 5 minutes after, gadolinium contrast injection, including the following flip angles: 1, 2, 4, 8, 12, 20, and 30°. All sequences in this series had a repetition time of 10 ms, echo time of 2.6 ms, pixel size of 187.5 μm x 187.5 μm x 200 μm, and 1 average. The scan time for the total image series (pre and post contrast injection combined) was ∼20 minutes. Gadolinium contrast (MultiHance) was injected as a bolus intravenously with a dose of 0.01 mmol diluted in saline at a molarity of 0.2mmol/mL.

### Generation of Gadolinium Concentration Maps

Separate pre-contrast and post-contrast T1 maps were calculated from each series of multiple-flip-angle 3D images using standard methods (Fram et al. 1987). Briefly, the MR signal magnitude as a function of flip angle *θ*_*n*_ was fit to the function

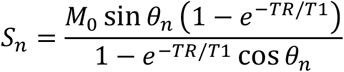

at each pixel containing nonzero signal, using the known value of the repetition time (TR = 10 ms) and sequence of flip angles *θ*_*n*_ = 1,2,4,8, 12,20, 30°. Deposited gadolinium concentration was then calculated at each brain pixel from the measured pre/post T1 change, using the known relaxivity of MultiHance (R=6.3 L/mmol/s) in the expression:

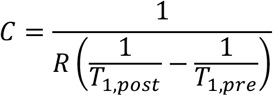

Physically impossible concentration values (those that were larger than the initial gadolinium concentration in the blood or significantly less than zero, usually occurring in image regions with low SNR) were excluded from further analysis.

### Segmentation of Lesions from MR Images

T2-SPACE images and gadolinium concentration maps were processed and analyzed in Horos DICOM viewer. T2-SPACE and gadolinium concentration maps were co-registered with the built-in feature within Horos or manually with custom MATLAB script. Manual segmentation of lesions were made with the freehand tool by outlining lesions in every coronal slice for each mouse and imaging timepoint within the study. Lesion volumes were calculated with the Horos “Compute Volume” feature and recorded. ROIs from T2-SPACE images were transposed onto co-registered gadolinium concentration maps. Total grayscale value within ROIs were recorded and summed across all slices for a given lesion (equivalent to gadolinium mass). Gadolinium concentration was calculated by dividing total gadolinium mass by computed lesion volume. Individual lesions were manually tracked across imaging time points using T2-SPACE images. The same anatomical location of a tracked lesion was validated in each imaging plane (coronal, axial, sagittal) across imaging timepoints. In some instances, distinct lesions from earlier time points had merged into one lesion at later time points. In other cases, lesions from earlier time points were no longer visible on images from later time points. De novo lesion formation was found in all animals in our study. If a lesion was obscured due to an imaging-related artifact, the lesion was excluded from analysis at the corresponding timepoint.

### Histology

After completion of the final imaging timepoint, mice were transcardially perfused with phosphate-buffered saline (PBS) and 4% PBS-buffered formaldehyde (EMS 15714). Brains were dissected, post-fixed, sequentially equilibrated in 10% and 30% sucrose and embedded in OCT (Andwin Scientific). Frozen brains were cryosectioned at 50-to 80-µm thickness for histology staining, or 25-µm thickness for immunohistochemistry (IHC). To perform histological staining of neurons and blood vessels in 50-µm thick sections, free floating sections were permeabilized overnight in PBS-buffered 0.5% Triton X-100 (Sigma-Aldrich 93443) at 4°C. Next, the sections were washed 3 times for 5 min with PBS, 1 mM CaCl_2_, and incubated in the red fluorescent Nissl stain (Neurotrace, Invitrogen N21482, diluted 1:100 in PBS, 1mM CaCl_2_) for 2 h at room temperature (RT). Following three washes with PBS, 1mM CaCl_2_, the sections were stained with Alexa Fluor 488-conjugated Isolectin GS-IB4 (Invitrogen 121411, diluted 1:100 in PBS, 1mM CaCl_2_) overnight at 4°C. After one final wash with PBS, 1mM CaCl_2_, the sections were mounted on microscopy slides with DAPI Fluoromount-G (SouthernBiotech, 0100-20) using Secure Seal Spacers (EMS 70327-20S).

### Immunohistochemistry

Thawed 25-μm sections were rehydrated with PBS, permeabilized in 0.5% Triton X-100 (Sigma-Aldrich 93443) in PBS for 1h at RT, and blocked with 1% bovine serum albumin (BSA, Jackson ImmunoResearch Labs, 001-000-161), 5% normal donkey serum (NDS, Jackson ImmunoResearch Labs, 017-000-121), and 0.5% Triton X-100 in PBS for 2 h at RT. The primary antibodies, including chicken anti-GFAP (1:200, Aves, GFAP5727980), rabbit anti-Iba1 (1:500, Wako Chemicals USA, 011-27991), goat anti-CD31 (1:20, R&D Systems, AF3628), were diluted in the blocking solution and incubated with mounted brain sections overnight at 4°C. After three 5-min washes with PBS, 0.5% Triton X-100, the sections were incubated with secondary antibodies, including donkey anti-chicken Alexa 488 (1:500, Jackson ImmunoResearch Labs, 703-546-155), donkey anti-rabbit Alexa 568 (1:500, Invitrogen A10042), donkey anti-goat Alexa 647 (1:500, Invitrogen A21447), diluted in the blocking solution at 1:500 for 2 h at RT. Following final washes, slides were mounted with ProLong Gold antifade reagent with DAPI (Invitrogen, P36935) and cover slipped with Fisherbrand Microscope Cover Glass (12-544-E) for confocal imaging.

### Confocal Microscopy

Stained sections were imaged with a Zeiss LSM 880 confocal microscope (Zeiss, Germany) using sequential scanning mode for DAPI, Alexa 488, 568 and 647 dyes. Montages of image stacks (1024 × 1024 pixels, 2 μm z-step), tiled in the x-y plane, were processed with Imaris 9.9 (Oxford Instruments) and analyzed with Fiji/ImageJ. Final images were adjusted with Adobe Photoshop and assembled in PowerPoint (Microsoft), or Adobe Illustrator (Adobe Creative Cloud).

### Segmentation of Lesions from IHC Images

Fluorescent images were analyzed in ImageJ for grayscale intensities in each channel: Iba1, GFAP, CD31, and DAPI. ROIs of lesion boundaries were manually drawn around the hypointense void space of lesions. To measure cell populations surrounding the lesions, the lesion boundary was expanded by either 50um or 100um, and the inside ROI was subtracted. Mean grayscale intensity of each of the three ROIs (inside lesion, 50um border, and 100um border) was averaged for all slices within the image stack in a given channel. Mean grayscale intensity for three reference ROIs drawn in nearby non-lesional locations of the brain tissue in each image were also averaged across the image stack in a given channel. Lesion associated mean grayscale intensities were normalized to the average reference mean grayscale intensity in the same image to account for variations of cell population expression in differing brain regions. These normalized mean grayscale intensities for Iba1, GFAP, CD31, and DAPI for each lesion were correlated with the same lesion’s volume and gadolinium concentration from the last imaging timepoint. Spearman’s correlation analysis was then performed with OriginPro’s Correlation Plot on the data matrix for 17 lesions and 14 measurements (3 lesion boundaries x 4 channels + lesion volume + lesion gadolinium concentration).

### Statistical analysis

The results were expressed as mean ± standard error of the mean (SEM). In all experiments, the statistical significance was set at P < 0.05. Calculations were performed using GraphPad Prism 8 statistical package software (San Diego, USA).

## Acknowledgments

This work was supported by funding from NIH R21NS116431 and grants from Focused Ultrasound Foundation, Be Brave For Life Foundation, and Alliance to Cure Cavernous Malformation to P.T.; NIH R01EB030409, R01EB030744, and R21NS118278 to R.J.P, NIH R01CA226899 to G.W.M., and AHA 830909 to D.G.F. We thank Dr. Kevin Whitehead for kindly providing the mouse strains used in this study. We are also grateful to Rene Jack Roy of the University of Virginia Molecular Imaging Core and Jeremy Gatesman of the University of Virginia Center for Comparative Medicine for assistance with MRI imaging and catheterization procedures, respectively.

