## Supplemental Material for "Understanding Lesion Progression in a Chronic Model of Cerebral Cavernous Malformations through Combined MRI and Histology"

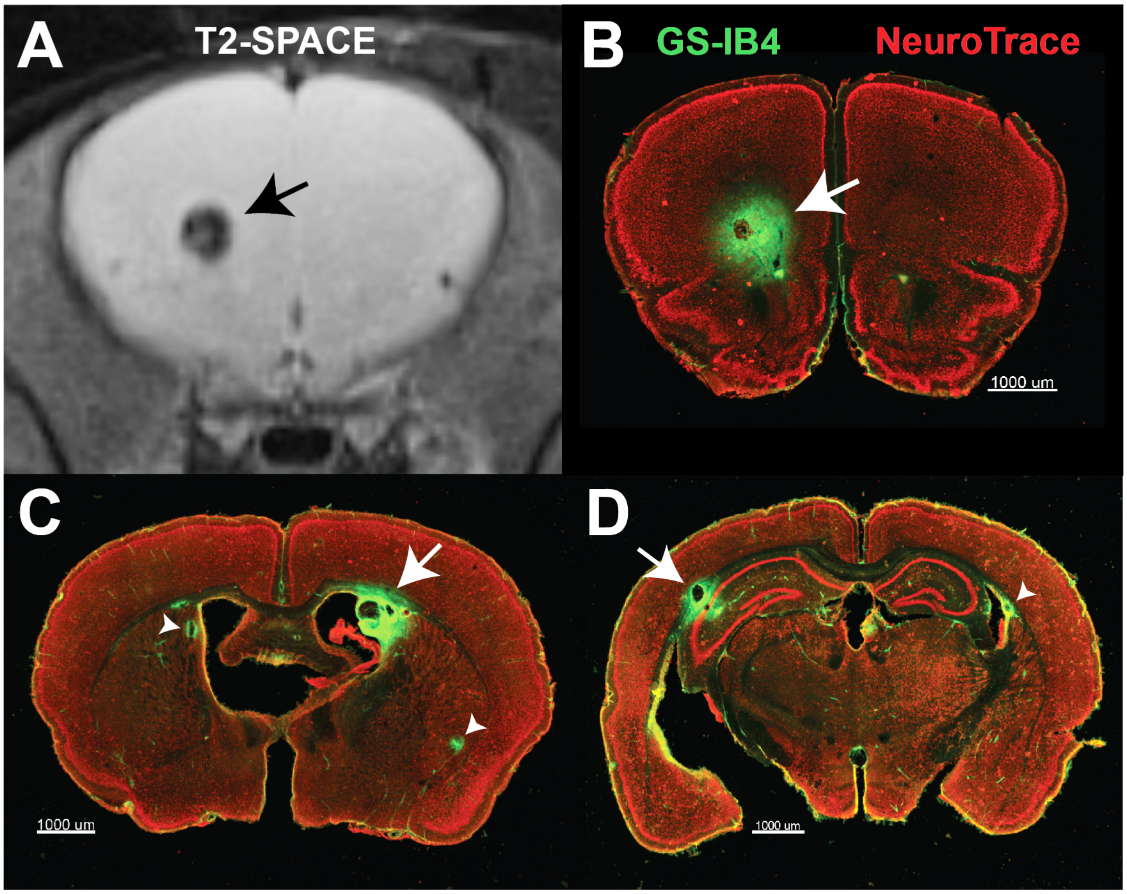


**Supplemental Figure 1.** The anatomical distribution of vascular lesions in the *Krit1* CCM model. **A, B,** Alignment of a T2-SPACE MRI image plane with the corresponding brain section labeled with fluorescent isolectin GS-IB4 (green) and Neuro-Trace Nissl stain (red). **C,** In this mouse model, cavernous lesions frequently occur in dorsal striatum near lateral ventricles (white arrow). **D,** CCM lesions are also commonly found in the subcortical white matter of the corpus callosum, often adjacent to the dorsal hippocampus. (white arrow, arrow head).

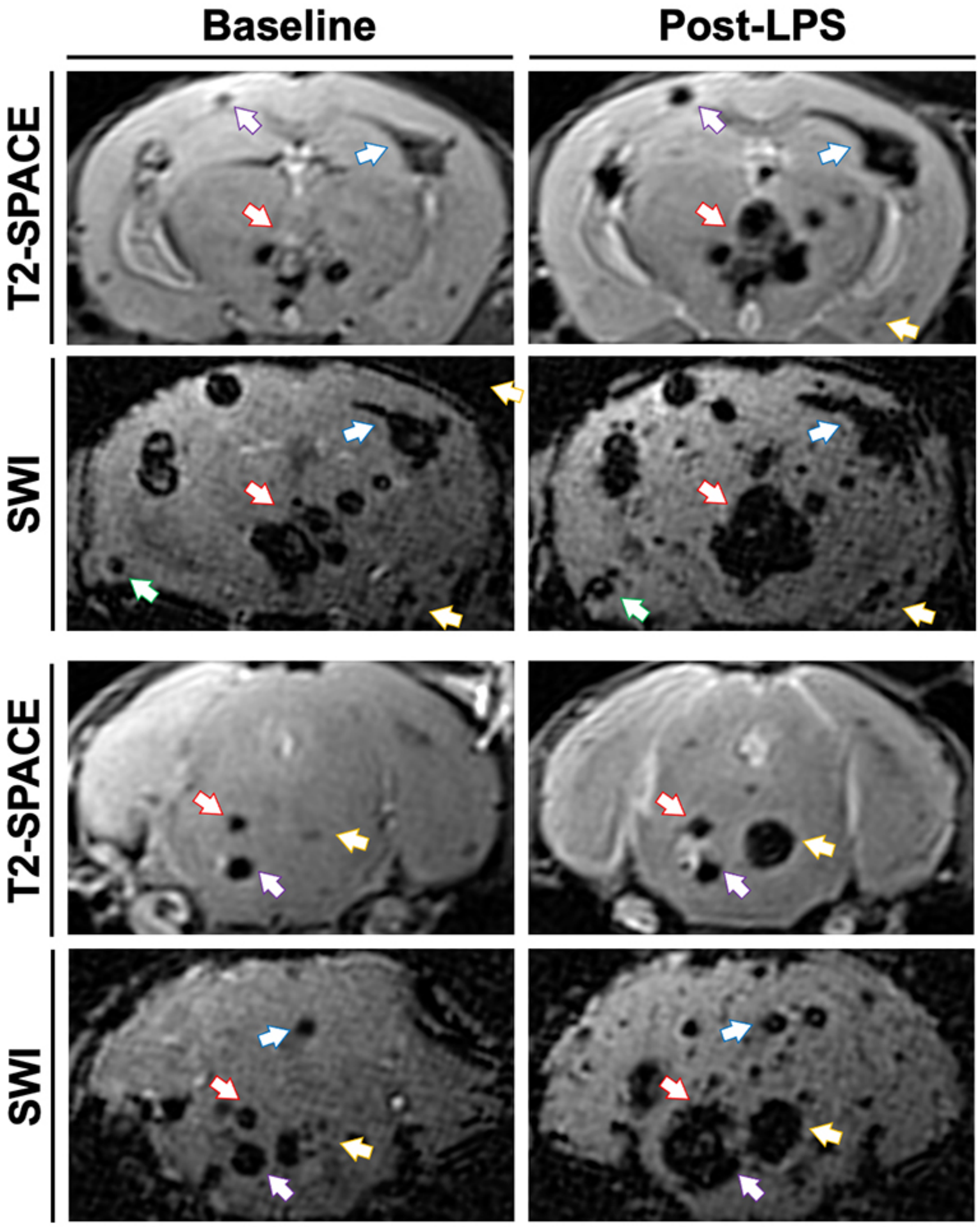


**Supplemental Figure 2.** Comparison of T2-SPACE and SWI sequence sensitivity before and after a sudden hemorrhagic episode. The *Pdgfb-CreERT*; *Krit1* $^{fl/null}$ mouse was imaged at Baseline and 12 h after acute mild neuroinflammation induced with LPS (0.25 mg/kg i.p.). Matching coronal planes of T2-SPACE and SWI are shown, demonstrating increased SWI sensitivity to CCM lesions. However, the exaggerated distortion of the lesion size is also apparent. Note the substantial increase in lesion volumes following neuroinflammation, accelerated due to fatal bleeding. Arrows identify the corresponding lesions between MRI sequences and inflammatory states.

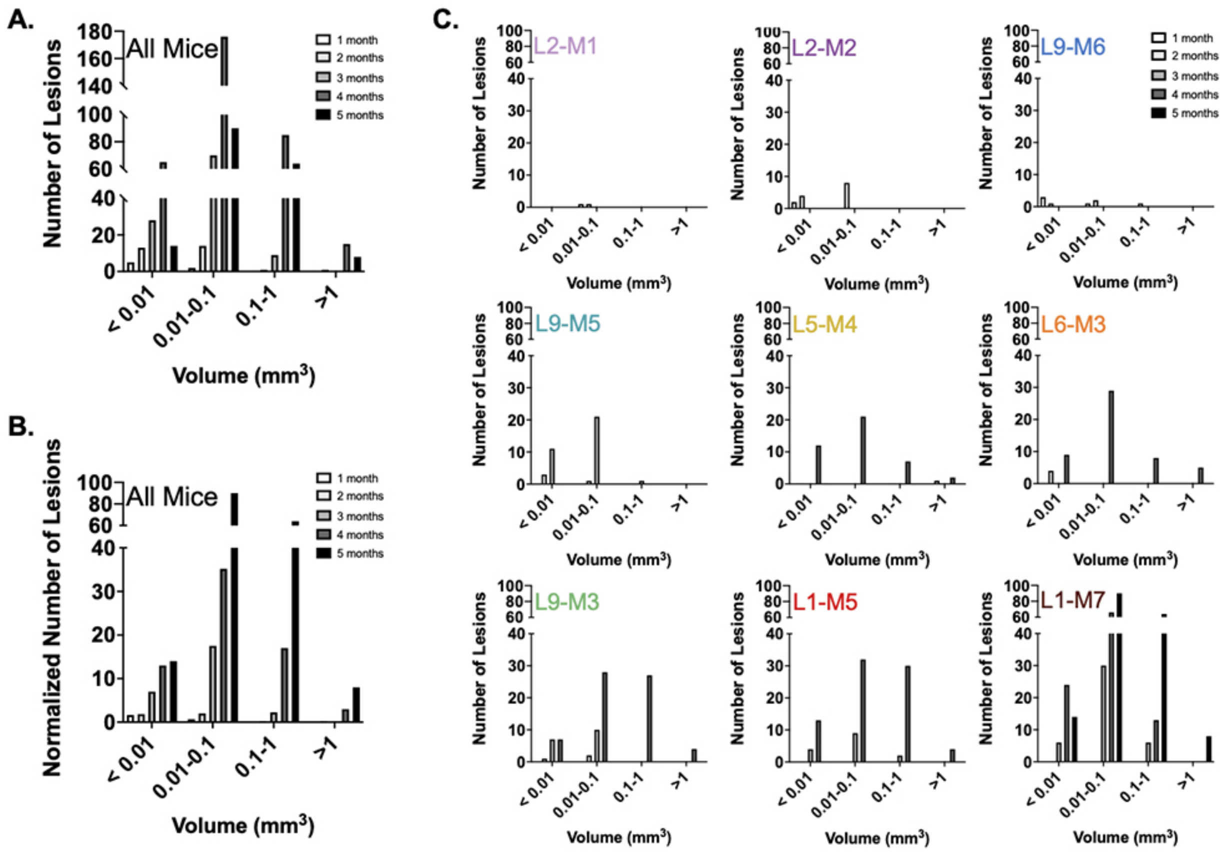


**Supplemental Figure 3.** Size distribution of lesions over time in our Krit1 chronic model. (A) Total number of lesions divided across four size categorizations for all mice in our cohort and each imaging time point. (B) Data from panel A normalized by the number of mice at each imaging time point. (C) Total number of lesions divided across four size categorizations for individual mice in our cohort and each imaging time point. Graph titles indicate unique mouse ID where L# denotes litter number and M# denotes arbitrary mouse number within litter.

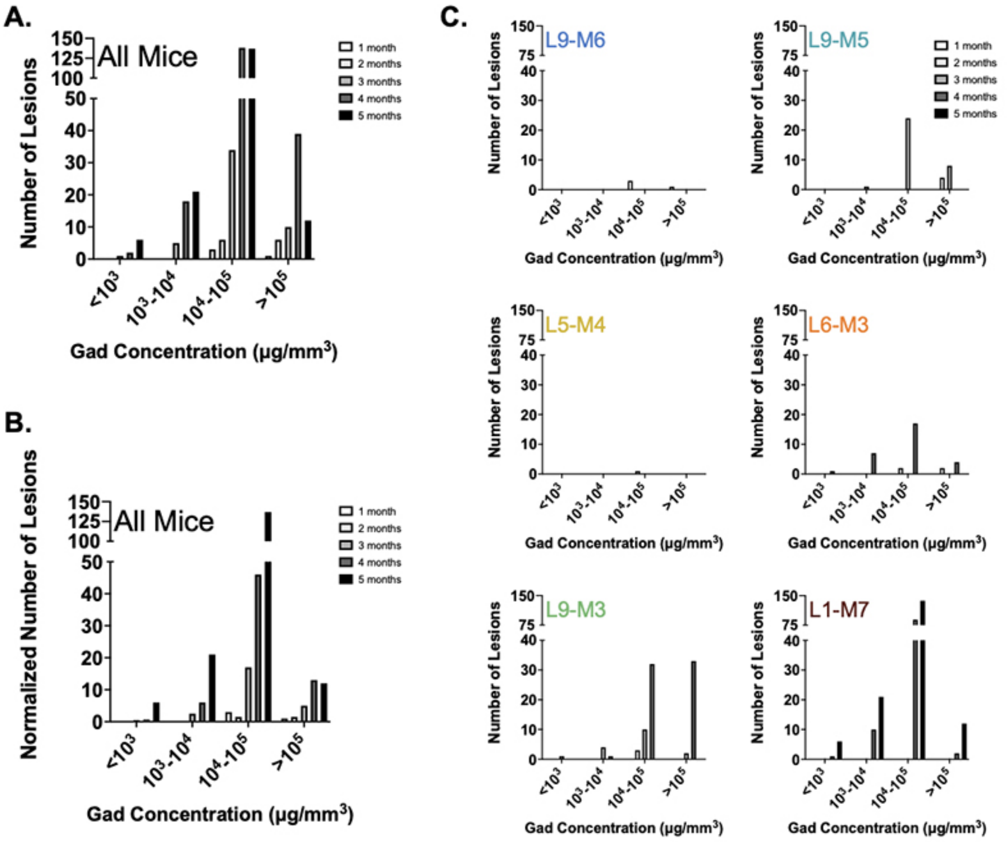


**Supplemental Figure 4.** Permeability distribution of lesions over time in our Krit1 chronic model. (A) Total number of lesions divided across four gadolinium concentration categorizations for all mice in our cohort and each imaging time point. (B) Data from panel A normalized by the number of mice at each imaging time point. (C) ) Total number of lesions divided across four gadolinium concentration categorizations for individual mice in our cohort and each imaging time point. Graph titles indicate unique mouse ID where L# denotes litter number and M# denotes arbitrary mouse number within litter.

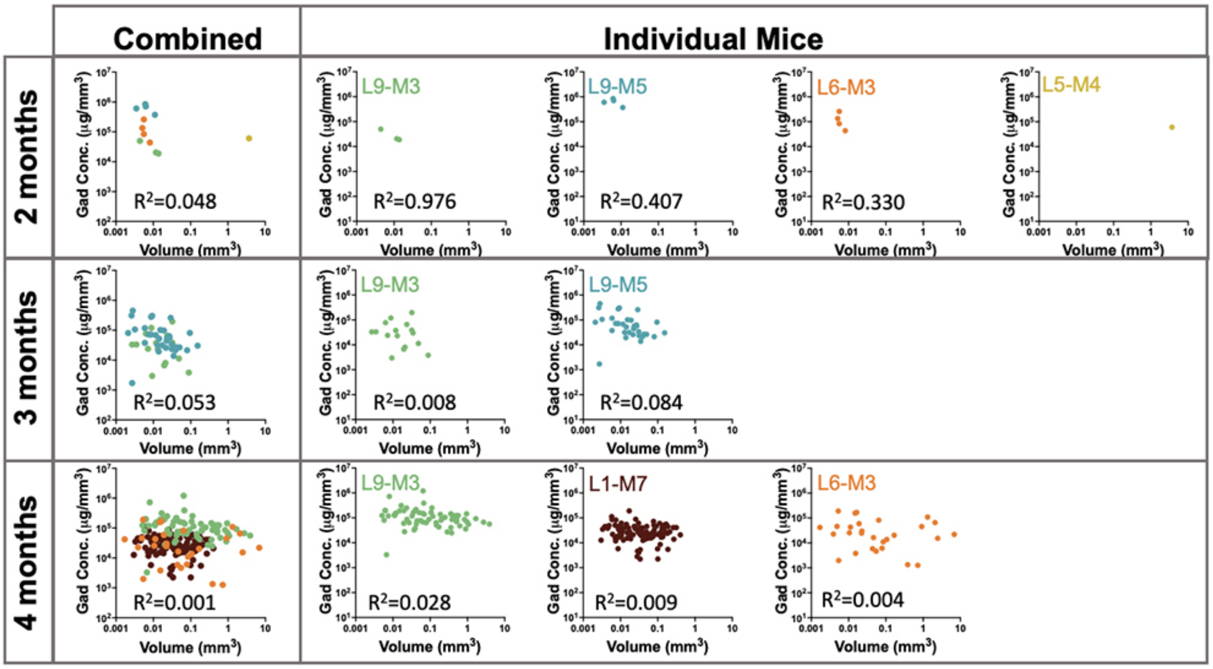


**Supplemental Figure 5.** Lesion permeability is poorly correlated with lesion volume. Graphs of gadolinium concentration as a function of lesion volume for individual lesions at each imaging timepoint. Data for all mice at 2 months, 3 months, and 4 months is shown on the left. Data for individual mice and corresponding coefficient of determination values are disaggregated on the right. Coefficient of determination values indicates poor correlation between gadolinium concentration and volume for each time point, suggesting highly heterogenous permeability of lesions across age in our chronic CCM model.

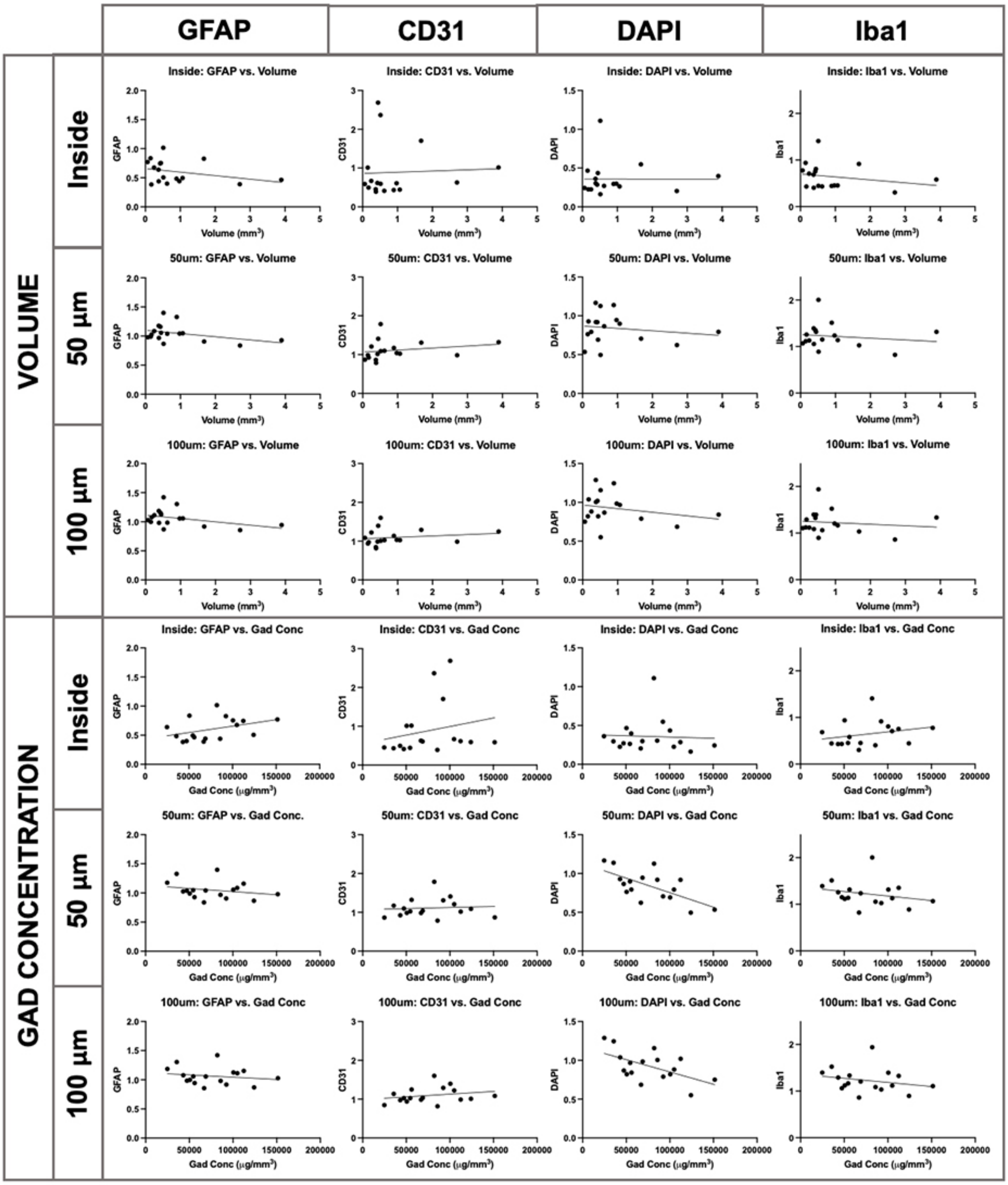


**Supplemental Figure 6.** Correlations between MR signatures and IHC signatures. Correlation plots for individual cell populations (astrocytes-GFAP, endothelial cells-CD31, cell nuclei-DAPI, and microglia-Iba1) and MR features (lesion volume and lesion permeability) at different locations around the lesion (inside lesion, 50μm border outside lesion, and 100μm border outside lesion). Dots indicate individual lesions (n=17). Linear regression fitted to each dataset.

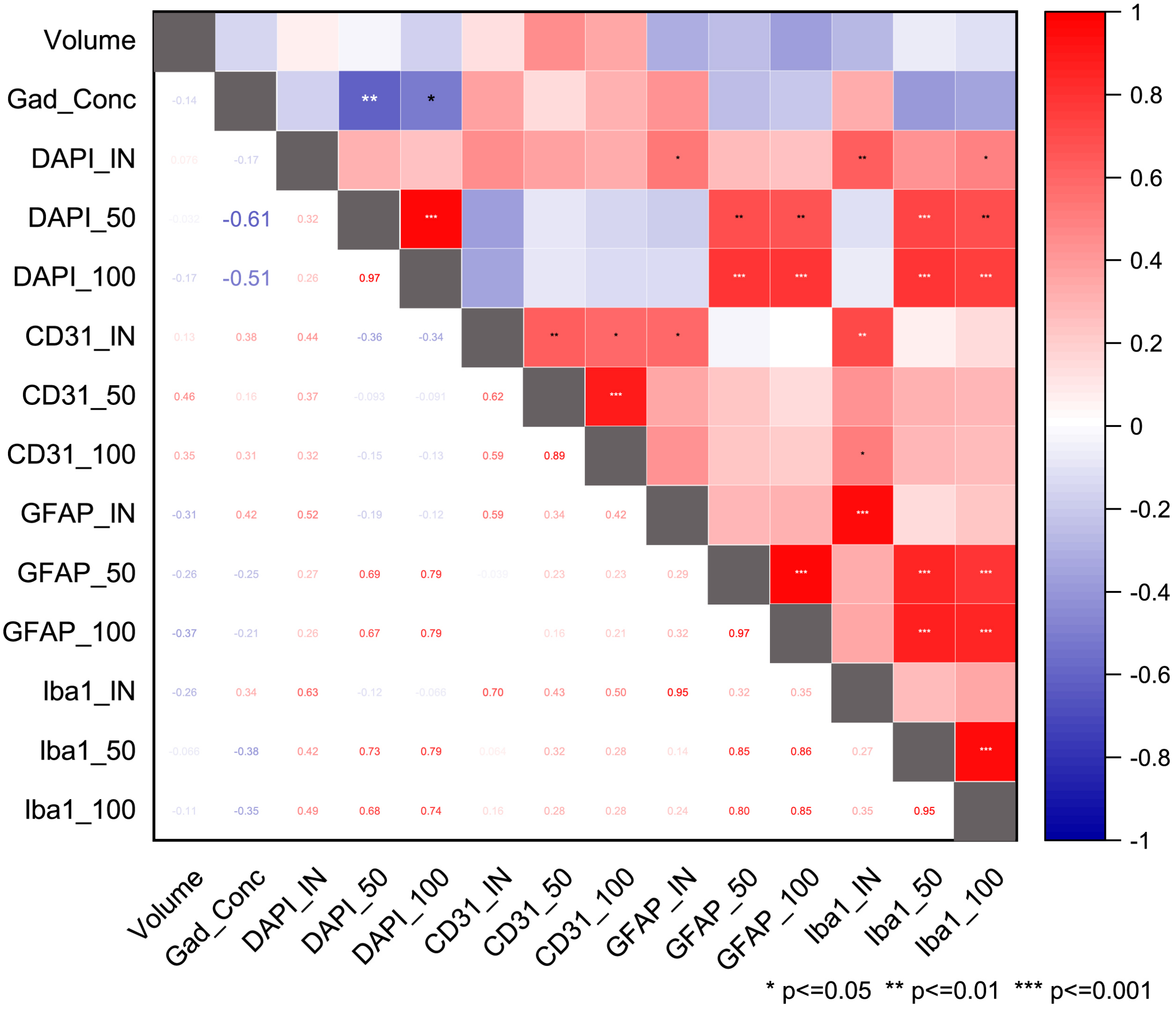


**Supplemental Figure 7.** Correlation Matrix Heatplot of 14 variables in 17 different CCM lesions in the L9-M3 animal. Two variables, Volume and Gad Concentration, were measured with MRI, the values of the remaining 12 variables were obtained with confocal image analysis of brain sections stained with IHC inside the lesions, as well as in 50- and 100-µm perimeters outside the lesions. Significant inverse correlations between MRI and IHC were found for gadolinium concentration and DAPI intensity for both 50-µm and 100-µm border perimeters ($\rho$ = - 0.61, p = 0.009; and $\rho$ = - 0.51, p = 0.036; respectively). The plot was generated with OriginPro 2020.

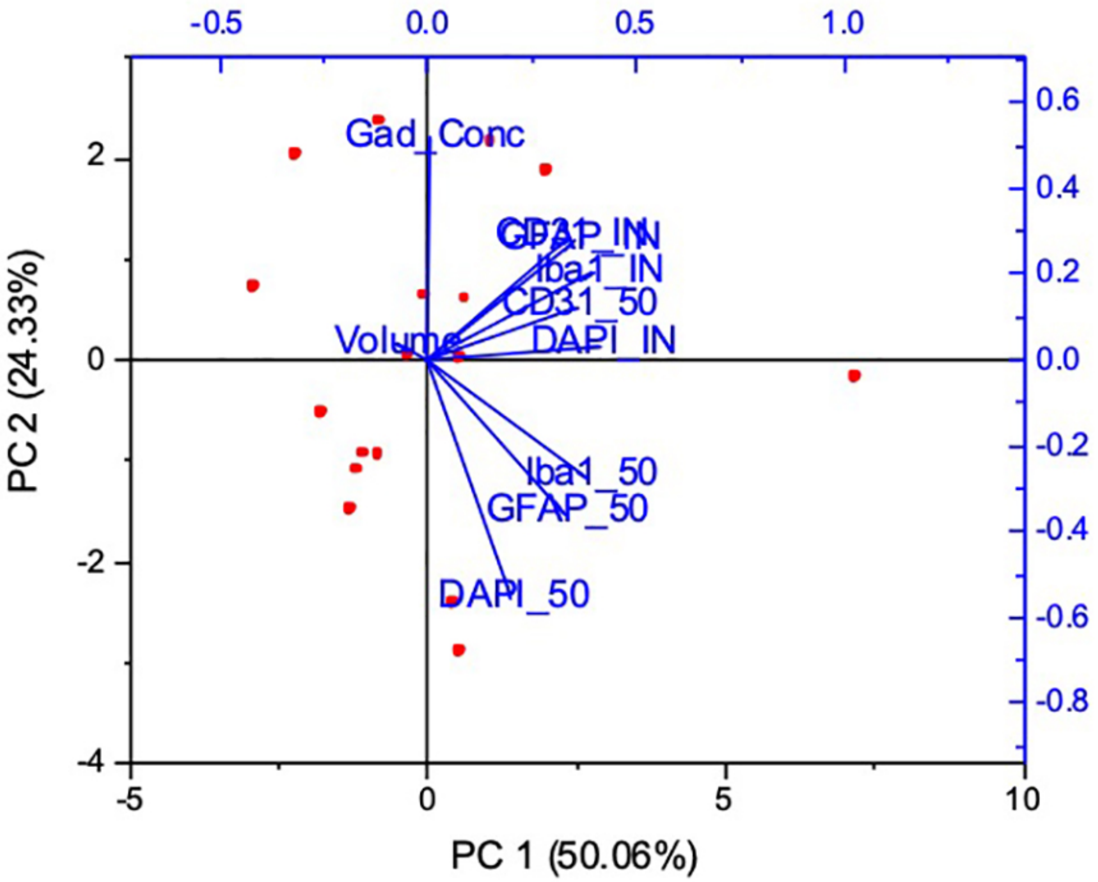


**Supplemental Figure 8.** Principal component analysis (PCA) of 10 variables in 17 different CCM lesions in the L9-M3 animal. (Including volume and gad. concentration, as well as the IHC values inside the lesions. Only the 50-µm perimeters were included in this analysis for the sake of clarity). PCA analysis corroborates the inverse correlation of gadolinium concentration and DAPI intensity outside the lesions, and reveals trends towards inverse correlation between gadolinium concentration and Iba1 and GFAP fluorescence intensity outside the lesions. The plot was generated with OriginPro 2020.

| Lesion | Volume | Gad_Conc | DAPI_IN | DAPI_50 | DAPI_100 | CD31_IN | CD31_50 | CD31_100 | GFAP_IN | GFAP_50 | GFAP_100 | Iba1_IN | Iba1_50 | Iba1_100 |
|---|---|---|---|---|---|---|---|---|---|---|---|---|---|---|
| 12 | 3.8942 | 56084.4333 | 0.39739129 | 0.793013312 | 0.841301184 | 1.015343379 | 1.321709253 | 1.248342177 | 0.464318626 | 0.928199992 | 0.943444281 | 0.580033178 | 1.315495626 | 1.333378936 |
| 16 | 0.9725 | 68803.0848 | 0.297844286 | 0.945526008 | 0.984488157 | 0.607504983 | 1.041377554 | 1.03322419 | 0.440994932 | 1.04330989 | 1.056577982 | 0.453377608 | 1.235461699 | 1.204742604 |
| 17 | 2.7031 | 66940.9197 | 0.204861705 | 0.622638739 | 0.685458344 | 0.628186609 | 0.982955394 | 0.987070813 | 0.389378661 | 0.836713562 | 0.855176704 | 0.303793737 | 0.821090259 | 0.860808859 |
| 18 | 1.6776 | 92340.248 | 0.547882669 | 0.705670112 | 0.789979156 | 1.702591788 | 1.308801057 | 1.292338025 | 0.826695824 | 0.905558174 | 0.91692457 | 0.916093035 | 1.02616352 | 1.034533021 |
| 19 | 0.5125 | 82025.3659 | 1.109236528 | 1.12710486 | 1.157201019 | 2.367267782 | 1.786515907 | 1.602337651 | 1.016410156 | 1.398065365 | 1.420632201 | 1.406452117 | 2.002086221 | 1.939948057 |
| 20 | 1.0659 | 54530.4438 | 0.26312907 | 0.89858737 | 0.96732064 | 0.44496801 | 1.023946598 | 1.026780424 | 0.496001559 | 1.047211018 | 1.057045842 | 0.453170135 | 1.138356628 | 1.165664988 |
| 30 | 0.6213 | 47234.8302 | 0.270757061 | 0.865818511 | 0.868262893 | 0.414967211 | 1.098258129 | 1.026857859 | 0.398852409 | 1.039804352 | 0.983979943 | 0.432314256 | 1.151532696 | 1.061633539 |
| 31 | 0.1706 | 43030.4807 | 0.226199119 | 0.928021675 | 1.037936966 | 0.49633306 | 0.924794108 | 0.972957703 | 0.384629348 | 1.025322037 | 1.077088337 | 0.430154292 | 1.256389644 | 1.290162734 |
| 32 | 0.3754 | 24813.5322 | 0.362126092 | 1.166441443 | 1.287928962 | 0.453880748 | 0.864685053 | 0.845853888 | 0.638901201 | 1.175176554 | 1.185035768 | 0.683475855 | 1.390881898 | 1.396404686 |
| 33 | 0.3832 | 85824.6347 | 0.304763751 | 0.919287784 | 1.004088445 | 0.391657715 | 0.786395762 | 0.815717693 | 0.438393894 | 0.96757882 | 0.980282254 | 0.402860512 | 1.054183823 | 1.085612833 |
| 34 | 0.894 | 35626.3982 | 0.295802019 | 1.137927448 | 1.246210492 | 0.433731044 | 1.170692197 | 1.136435801 | 0.483419921 | 1.327219625 | 1.304772727 | 0.443061046 | 1.512346836 | 1.524060222 |
| 43 | 0.424 | 112400.943 | 0.285584303 | 0.918074989 | 1.019386527 | 0.616323182 | 1.015632842 | 0.988804731 | 0.745158523 | 1.159673947 | 1.152775514 | 0.752698686 | 1.353873897 | 1.326104034 |
| 44 | 0.442 | 100450.226 | 0.435529058 | 0.692213627 | 0.820822228 | 2.685201114 | 1.408309008 | 1.397247039 | 0.75415504 | 1.056234735 | 1.125533394 | 0.807036337 | 1.316149344 | 1.395134376 |
| 45 | 0.1478 | 50663.0582 | 0.466523555 | 0.762592968 | 0.820315419 | 1.011510244 | 0.98743172 | 0.939195821 | 0.836290649 | 0.989456106 | 0.99701947 | 0.937812807 | 1.114896027 | 1.122397233 |
| 46 | 0.5158 | 124189.608 | 0.162594569 | 0.49567473 | 0.550319294 | 0.593272696 | 1.087941829 | 1.007667632 | 0.505722136 | 0.866029998 | 0.86956832 | 0.444563928 | 0.888560425 | 0.896209433 |
| 47 | 0.0632 | 151503.165 | 0.243078743 | 0.534685677 | 0.751097191 | 0.588777686 | 0.868744458 | 1.08461039 | 0.771367845 | 0.979777542 | 1.027360352 | 0.776609943 | 1.067659463 | 1.106251985 |
| 50 | 0.2495 | 104917.836 | 0.225765379 | 0.792745439 | 0.881565478 | 0.667507802 | 1.208254576 | 1.223397011 | 0.675876461 | 1.085660367 | 1.112661988 | 0.707738575 | 1.129422826 | 1.117259334 |

**Supplemental Table 1**. MRI and IHC measurements of 17 lesions obtained from animal L9-M3.
